# Bedsect: An integrated web server application to perform intersection, visualization and functional annotation of genomic regions from multiple datasets

**DOI:** 10.1101/481333

**Authors:** Gyan Prakash Mishra, Arup Ghosh, Atimukta Jha, Sunil Kumar Raghav

## Abstract

A large number of genomic regions captured during next generation sequencing (NGS) data analyses are generally overlapped to answer a variety of biological questions. Though several command-line tools are available to perform such analysis, there is lack of a comprehensive web server application to perform the genomic region intersections from multiple datasets, to generate a plot for the subsets of the overlapped regions and to perform integrated functional annotation. To address this gap, we have developed a user-friendly integrated web server application i.e. Bedsect, where users can upload genomic regions from multiple datasets without any file number limitation and perform intersection analysis along with visualization of the intersection regions as UpSet and Correlation plot using integrated Shiny application. Bedsect also integrates GREAT for functional annotation, gene ontology, and biological pathways enrichment analysis from identified unique as well as intersected genomic regions. These genomic regions can be further uploaded in the UCSC genome browser for visualization of the results as custom tracks directly from the tool. Bedsect is available at http://imgsb.org/bedsect/.

## Introduction

With the decline in cost, Next Generation Sequencing technology is becoming a popular method to identify genomic regions of various interest such as transcription factor binding regions, accessible chromatin regions, regions with histone modification, methylated regions, frequently interacting regions etc. These binding events/chromosomal locations are generally stored in BED (browser extensible data) [1] format files. Overlapping of these regions is generally used to answer important questions to understand gene regulation, chromatin architecture etc. Investigators mostly overlap genomic regions generated in different analyses to make important biological interpretations and conclusions, for example, to identify genomic HOTSPOTS bound by multiple TFs, *de novo* DNA motif predictions at these sites, comparing DNaseI-hypersensitive sites (DHS) across different cells types, Conserved non-coding Elements, etc. Many command-line based tools such as BEDtools [2], BEDOPS [3], Intervene [4] are available to overlap BED files. However, these tools require some knowledge of linux command-line to carry out such analysis. Several other R packages such as ChIPpeakAnnon [5], GUI based and web server tools such as PeakAnalyzer [6], PAVIS [7], LOLA [8] are also available to perform functional annotation of the genomic regions. The above-mentioned tools undoubtedly are very useful for performing genomic region intersection analysis or functional annotation. However, the above-mentioned tools come with specific functionalities of either only region overlap calculation, only visualization or functional annotation. To overcome these limitations we developed comprehensive web server application Bedsect, that provides an integrated platform for genomic region intersection without any number limitation for datasets to upload for overlap, visualization of intersection regions as UpSet plot [9] and correlation plot using an integrated Shiny application (ShinyApp). Moreover, it also generates individual BED files for unique as well as overlapping regions between the datasets that can be downloaded from the result page generated by the tool. Further ShinyApp provides user to generate high-quality images for reporting. To perform functional annotation of these regions, integrated tool Genomic Regions Enrichment of Annotations Tool (GREAT) API [10] can be used that does annotation in terms of distribution of genomic regions near to transcription start site (TSS), GO enrichment, pathway enrichment, and enrichment against many other associated databases. Users can also use links within the tool itself to upload intersected and unique genomic regions in University of California, Santa Cruz (UCSC) genome browser for quick visualization of the results [11].

One of the important aspects in functional genomics is to identify genomic regions bound by a diverse set of transcription factor (TF) and associated with histone modifications (either at promoter or enhancer regions) that marks chromatin structure changes [12, 13]. To identify the distribution of such genomic regions near TSS, and functionally annotate genes regulated by cis-regulatory elements bound by multiple regulatory factors, integration of publicly available web server tools such as UCSC genome browser and GREAT would be of great advantage. [10]. PCR / ChIP-qPCR could be performed to validate the predicted binding of different transcription factors, once such regions are determined by Bedsect. Region-specific PCR primers are required for validation experiments. Therefore UCSC API generated hyperlinks provided by our tool to visualize the genomic regions as custom tracks in genome browser would be quite helpful. As there is no such easy to use and efficient user-friendly web server tool to overlap multiple BED files, Bedsect would be of great advantage to a broad audience working in the field of functional genomics.

## Design and Implementation

The core of this tool utilizes Perl, R, Shiny server and the front end of the web server written using PHP5.6, MySQL 14.14; Dist 5.7.19 and Javascript 1.8. User uploaded input files (BED format) on the web server is processed to find overlap using the multiIntersectBed program of BEDtools. Using a parser written in Perl language, the output file is parsed to extract overlapping regions between different datasets. The output generated from multiIntersectBed contains total genomic regions across all the datasets and its presence or absence across datasets represented in terms of 0 and 1 (0 for absence and 1 for presence) in a binary matrix table. The matrix table is used to generate results as UpSet and correlation plots (Fig 1). To identify the similarity between different datasets, it calculates pairwise Jaccard index between datasets from the obtained binary matrix and generates a correlation plot. Further, to carry out functional annotation of the intersection regions, we have integrated direct access to GREAT server that annotates genomic regions to target genes and calculates statistical enrichment for the association of genomic regions (Fig 2B). Default setting of tool that annotates genomic regions to genes based on distance to TSS (proximal: 5kb upstream to 1kb downstream; distal; up to 1000kb) has been implemented, but if users intend to use other parameters, BED files of intersecting or unique regions can be downloaded and uploaded in GREAT with other settings. Our tool also provides an option as a hyperlink to submit unique or intersecting genomic regions as a custom track to visualize in UCSC genome browser (Fig. 2B)

**Fig. 1:**
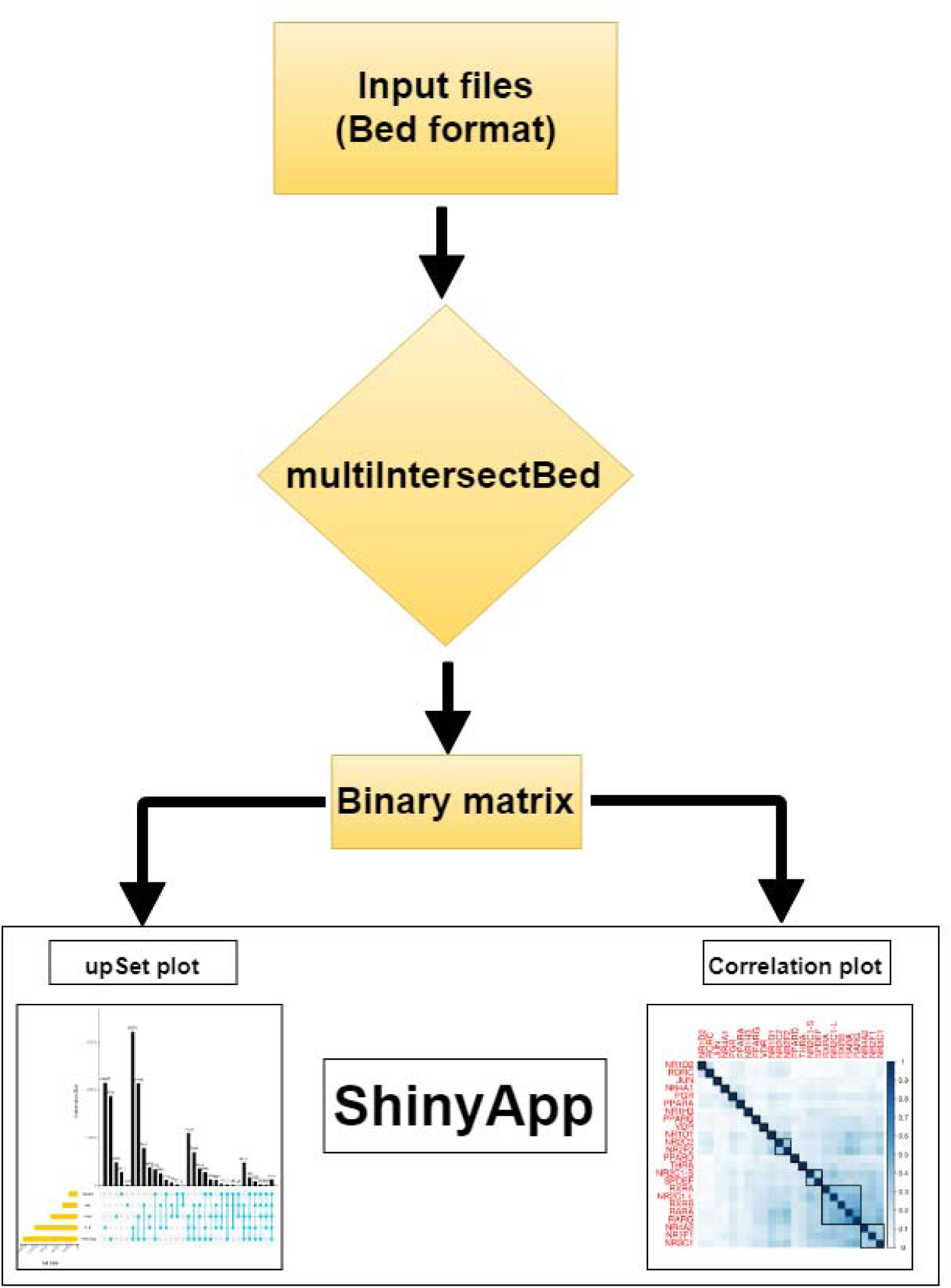
Flowchart outlining workflow of Bedsect. Input files in the BED format is analysed to find overlap using multiIntersectBed of BEDtools with default options and binary output is processed to generate UpSet plot and correlation heatmap using ShinyApp.

**Fig. 2:**
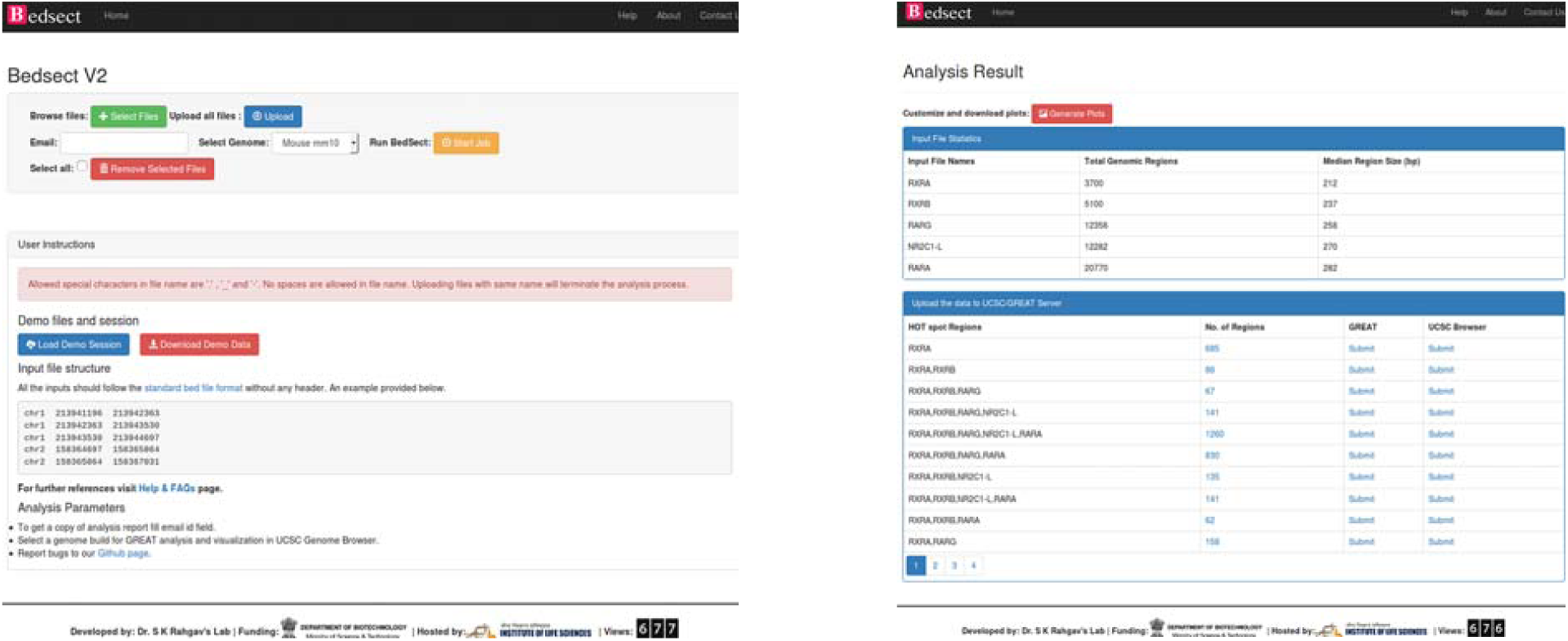
(A) Snapshot of the homepage of the web server. (B) Result page after uploading BED files.

## Web server

The homepage of Bedsect features an input form with a set of fields such as browse files, genome, user email id, and bed file upload (Fig. 2A). Selecting appropriate genome version is required to perform functional annotation and genome browser visualization. Depending upon the availability of server resources, number and size of files, the analysis may take several minutes, so filling up the email id is mandatory and will be used to notify links to Bedsect analyzed results. After every successful submission, the user will be redirected to results page after 5 seconds, depending upon the status of analysis specific message will be displayed. For overlapping two or more files, the user should select the BED format files having extension “.bed” using “Select Files” tab and “Upload” tab to upload files. To remove any uploaded files, the user can also select “Remove Selected Files” tab to remove the uploaded files. After a successful upload, “Start Job” tab is used to perform the analysis. Once submitted user will receive an email for job submission as well as for job completion. After completion of the analysis, a result page is loaded that includes metadata providing total genomic regions and the median value of genomic region size (bp). Moreover upon selection of any genome, while submitting the files, result page will give an option in tabular format to carry out further analysis of intersecting regions using GREAT and UCSC (Fig 2B). Following customized and easy to interpret publication-ready plots can be generated using integrated ShinyApp.

## UpSet plot

One of the best alternative to visualize comparison of datasets other than Venn diagram is an UpSet plot. After a certain threshold interpretation of Venn diagram becomes difficult so our Bedsect tool provides an easy to interpret UpSet plots. The ShinyApp also provides various options to customize the generated plot (Fig 3).

**Fig. 3:**
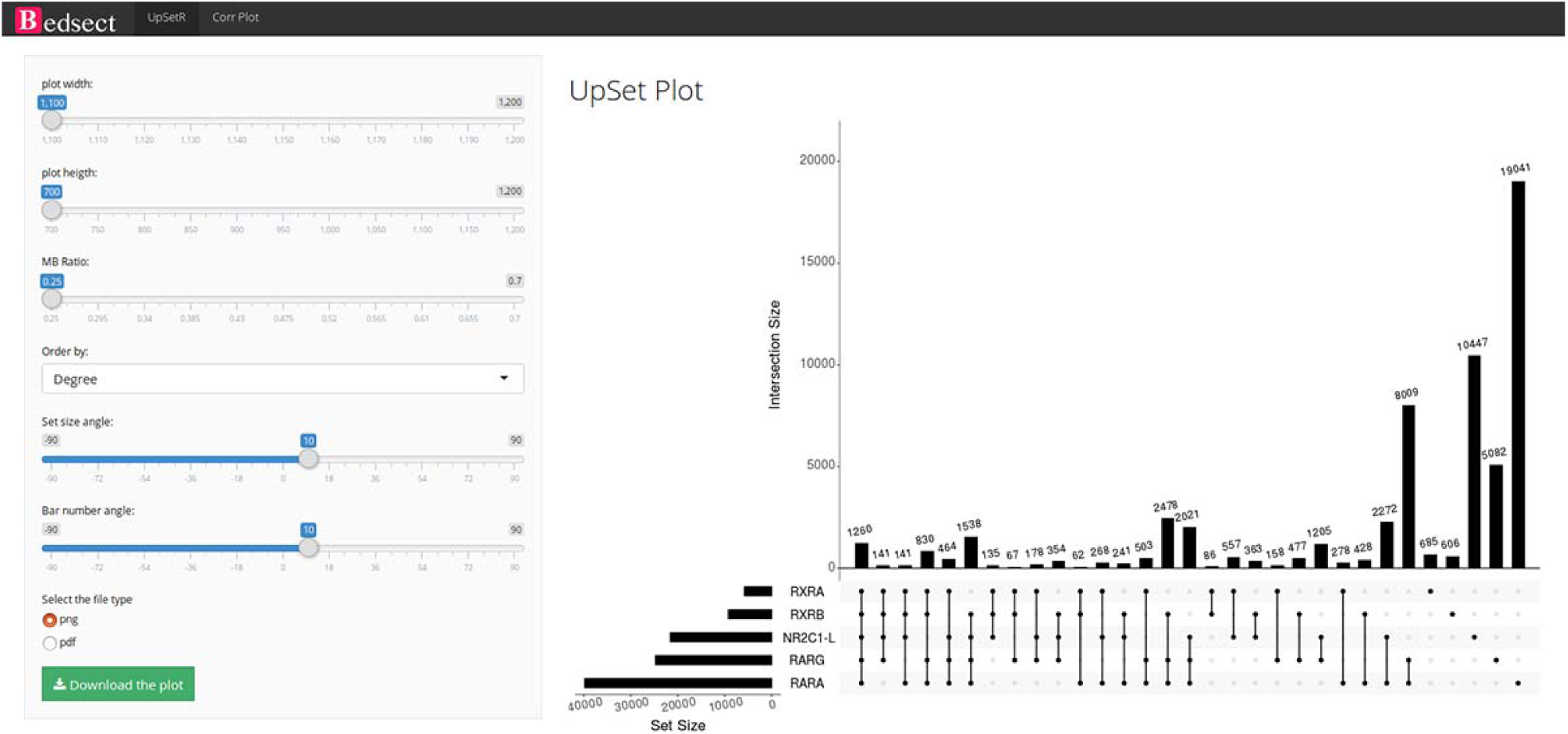
UpSet plot representation of intersection regions.

## Correlation plot

To calculate the correlation between different datasets, we implemented Jaccard function of an R package SuperExacttest [14] which calculates Jaccard statistics based on a total number of overlapping regions between datasets. The estimated values are then plotted as correlation heatmap using R corrplot package [15]. In addition, ShinyApp provides the users various option to customize the plot (Fig. 4).

**Fig. 4:**
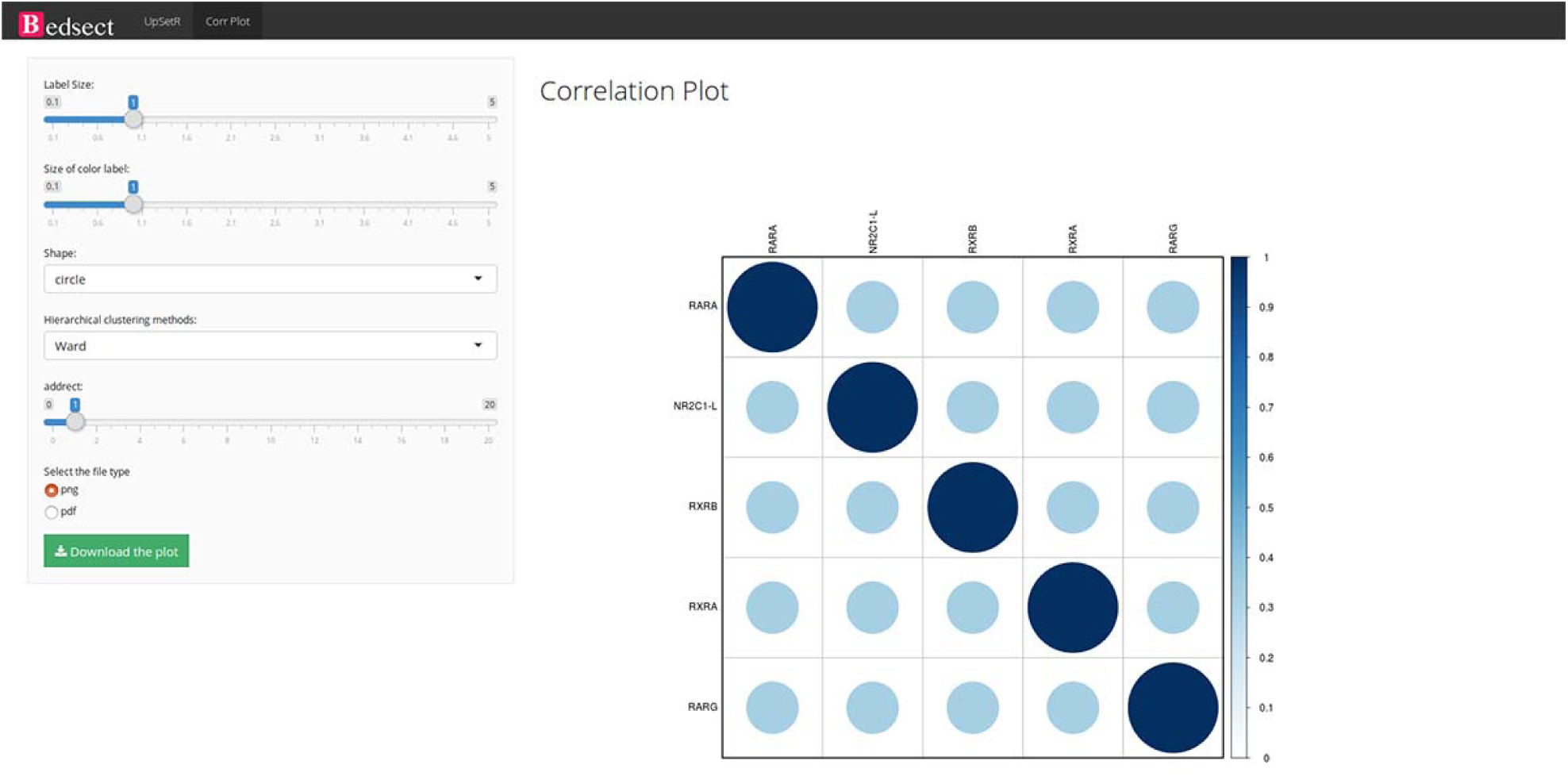
Correlation plot representing similarity based on Jaccard indices (0 to 1).

## Functional annotation and visualization in UCSC browser

To perform functional annotation and track visualization, currently, we have provided Mouse (mm10 and mm9), Human (hg19) and Zebrafish (zv9) genome. Upon selection of genome build at home page, the result page provides a hyperlink option to submit the unique and intersecting regions to GREAT server and visualization in UCSC browser (Fig 5).

**Fig. 5:**
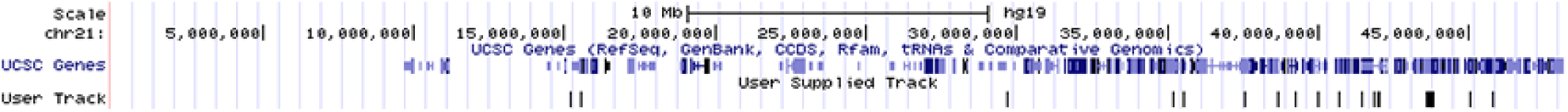
Snapshot of UCSC genome browser showing uploaded user track.

## Case Study

To demonstrate the utility of our tool, we studied the genome-wide binding of multiple transcription factors. Regions co-bound by multiple regulatory factors or histone modifications are considered HOTSPOTS that strongly regulate gene transcription. We downloaded online available datasets from a published study using MCF7 breast cancer cell line, where 24 nuclear receptors (NR) bindings were profiled using NGS (GSE41995) [12]. The genome version of all the files was lifted from hg18 to hg19 using liftOver tool from UCSC utilities. All the converted files are available at https://github.com/sraghav-lab/Bedsect/tree/master/test_data. It has been reported that the HOT regions (regions occupied by multiple transcription factors) plays an important in cancer development [12]. To identify these regions, here we implemented our comprehensive web server application. We overlapped 24 NR peak files in BED format. To identify factors that have high genome-wide binding similarity, we generated correlation plot using integrated ShinyApp by clicking on “Generate plots” tab. To identify correlation, it calculates pairwise Jaccard index to develop correlation plot using R corrplot package. We demonstrated that RARα, RARγ, RXRα, RXRβ and NR2C1-L bindings are highly similar based on genome-wide occupancy of all these factors (Fig. 6). Study published by Kittler et al has also shown using network analysis that RARα, RARβ, RXRα and RXRβ regulates genes associated with breast cancer development. Next we uploaded only these five datasets to identify regions co-bound by only these five transcription factors. In addition, to identify the distribution of the overlapped regions near TSS, we submitted the identified regions (n=1262) (Fig. 7) co-bound by these five trans-factors to GREAT. We found that majority of the bindings are present in far-distal regions (>5kb to TSS) (Fig. 8A). The gene annotation plot showed that 12 regions were not annotated to any gene, 156 regions annotated to one gene and each of the 1092 genomic regions annotated to two genes (Fig 8B). Furthermore, to predict the functional role of the genes annotated to intersected genomic regions or HOT regions, we looked into the enriched pathway against MSigDB. The enriched terms indicated the association of annotated genes to breast cancer, gastrointestinal tumor, epithelial to mesenchymal transition (Fig. 8C). Overall analysis of the presented datasets suggested that our Bedsect tool is quite efficient and powerful to identify regions intersecting between different datasets and to help in quick prediction of the functional importance of the intersected genomic regions.

**Fig. 6:**
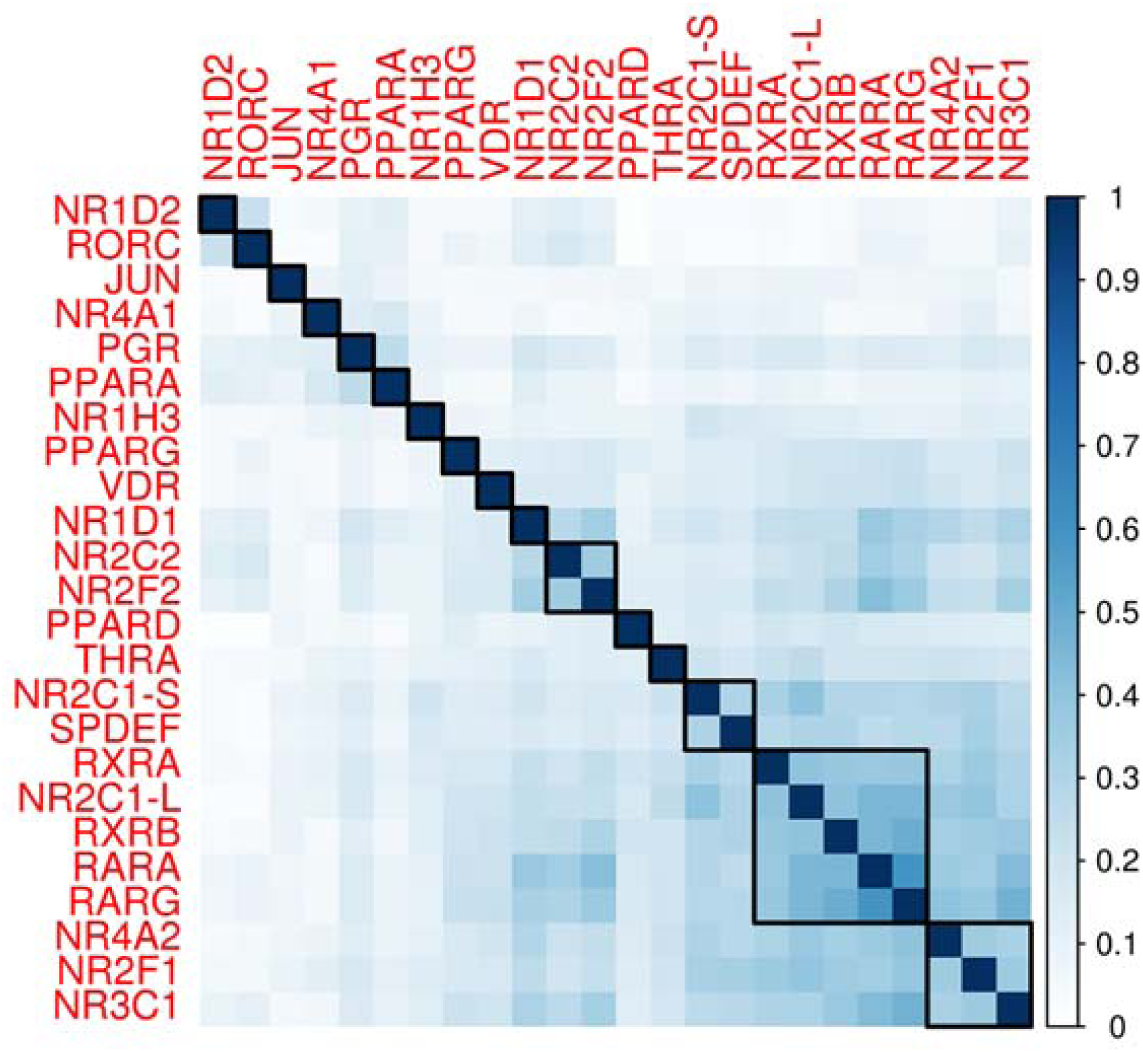
Heatmap of correlation of 24 NR based on Jaccard index.

**Fig. 7:** UpSet plot representing the number of subsets of intersection between five factors.

**Fig. 8:**
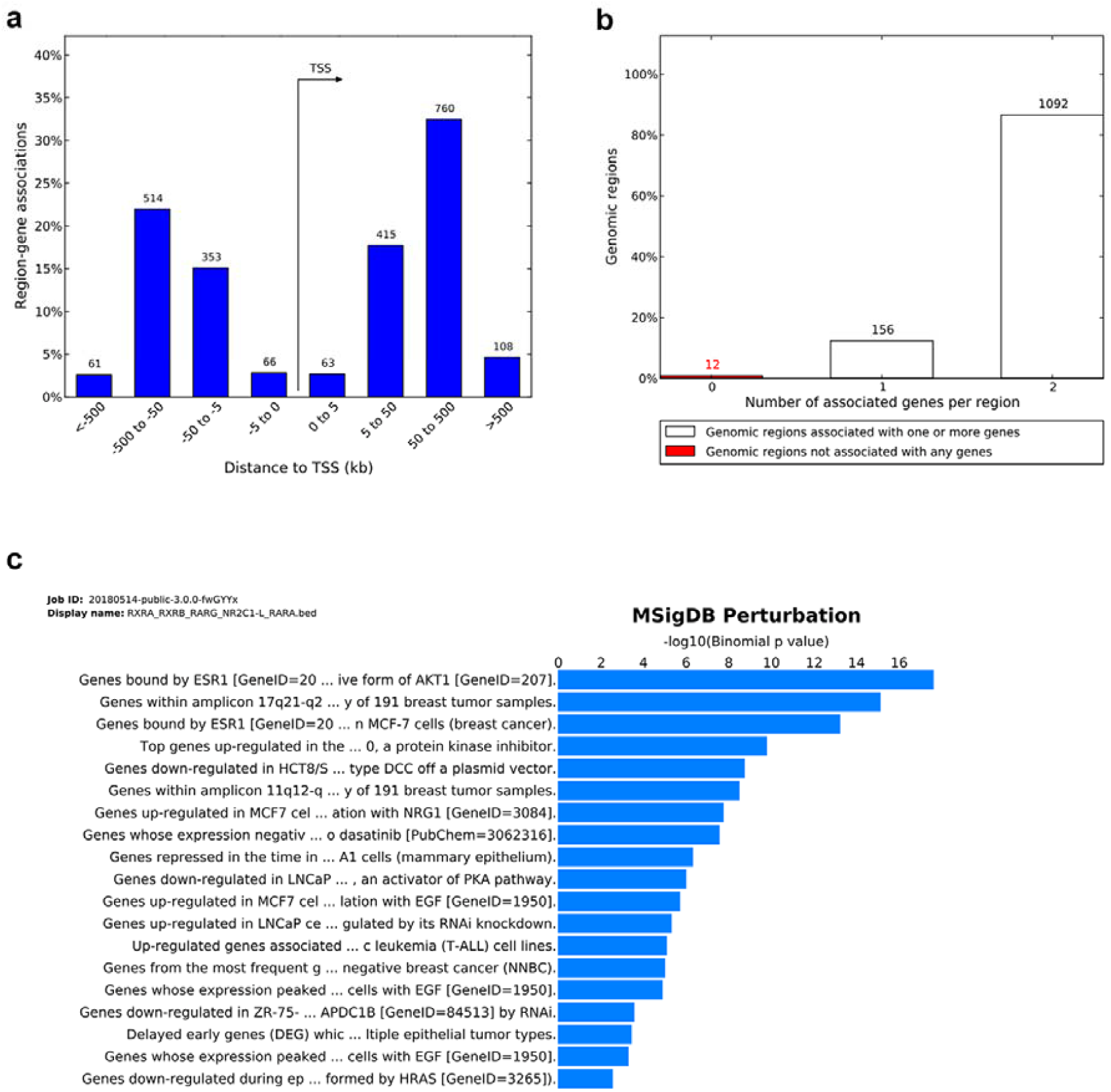
(A) Genomic regions distribution near TSS. (B) Number of genes annotated to genomic regions. (C) Enriched pathway against MSigDB obtained from the GREAT analysis.

## Comparison with other publically available tools

To show advantage and usefulness of our tool Bedsect, we compared it with various publically available tools based on various attributes. Table 1 shows the comparison of Bedsect with all other currently available tools.

**Table 1:**
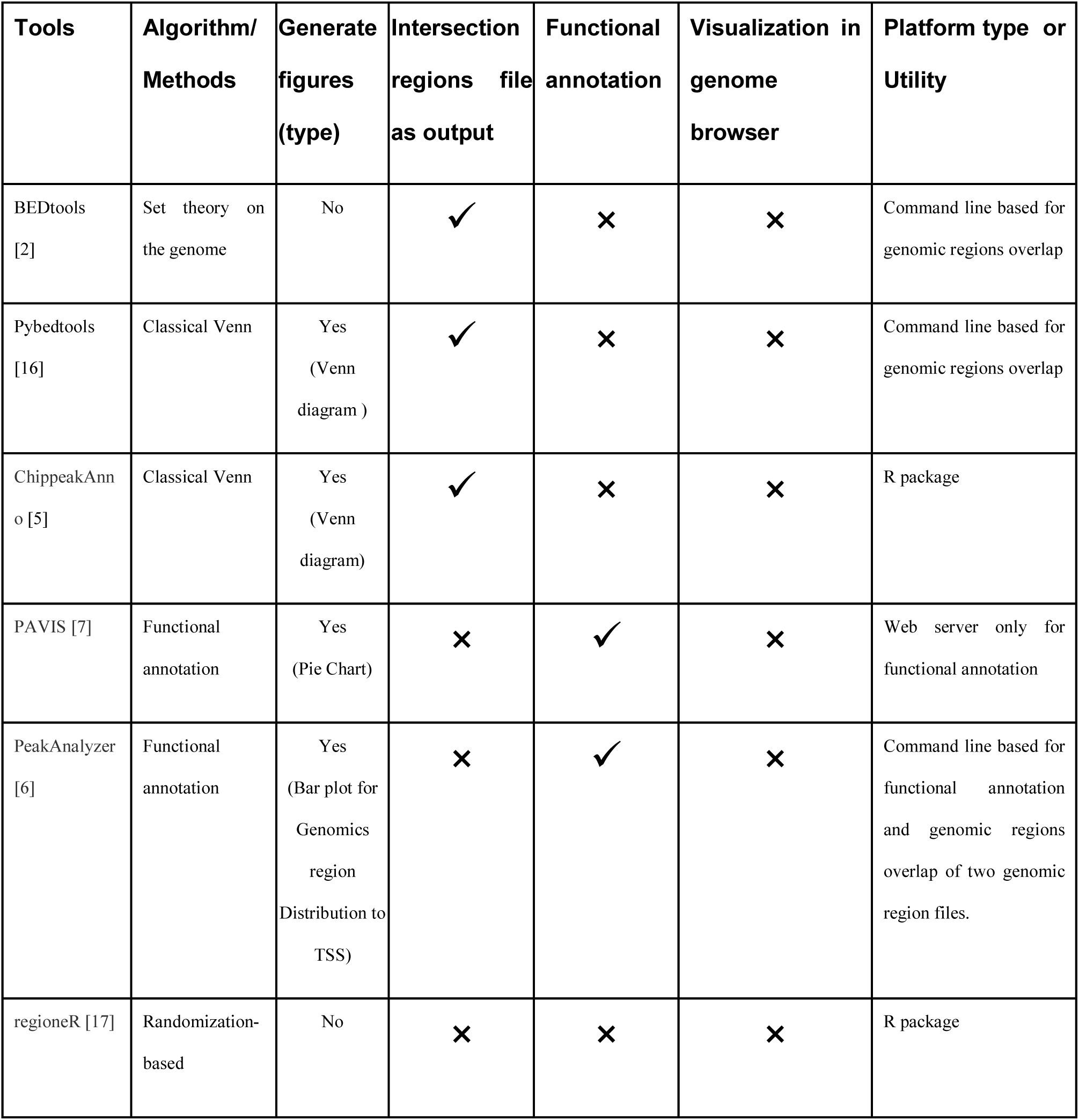
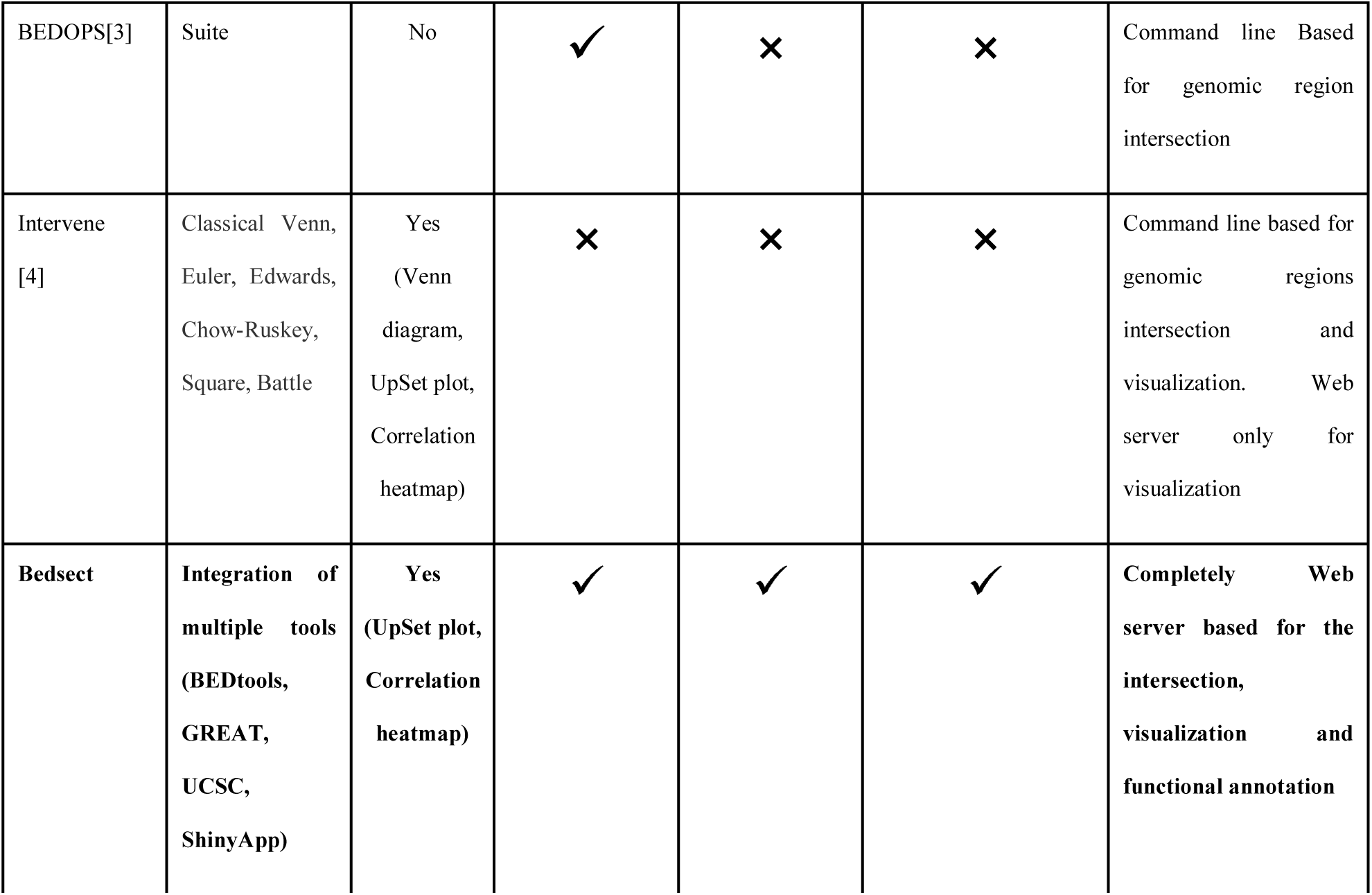
Comparison of Bedsect with other publically available tools

## Availability

The tool is available at http://imgsb.org/bedsect/ and source code is available at Github (https://github.com/sraghav-lab/Bedsect)

## Acknowledgments

We are grateful to the developer of a very efficient tool BEDTOOLS and other tools that we have used to develop our web server tool.

